# High-throughput screening for class I peptide MHC binding via yeast surface display

**DOI:** 10.1101/2025.05.24.655874

**Authors:** Patrick V. Holec, Kathryn C. Breuckman, Owen Leddy, Forest M. White, Bryan D. Bryson, Michael E. Birnbaum

## Abstract

T cells rely on short peptides presented by highly polymorphic major histocompatibility complexes (MHCs) to selectively initiate adaptive immune responses. Despite its importance, few techniques can systemically evaluate stable peptide presentation across diverse MHC alleles. Here, we describe a yeast display pipeline that can be deployed to rapidly screen proteomic space to identify class I pMHC binders across many alleles. Through this, we capture unique biological phenomena such as interference with peptide presentation via type IV drug-induced hypersensitivity. We apply this approach to multiple pathogen proteomes (Mtb Type 7S substrates, SARS-CoV-2, Dengue, and Zika) to create a high-resolution catalog of potential T cell antigens. Altogether, this platform acts as a flexible tool to generate large unbiased datasets for class I peptide presentation at a speed and scale competitive with the biological systems they represent.

## Introduction

Adaptive immune recognition depends upon the presentation of antigens by Major Histocompatibility Complexes (MHCs)^1^. MHCs are membrane-bound proteins that display short peptide fragments to convey information of cell state to T cells. MHCs are divided into two classes that dictate their biological function. Class I MHCs are expressed on every nucleated cell in the body and primarily display peptides derived from the host cell, while class II MHC are expressed on professional antigen presenting cells and primarily display peptides derived from the extracellular milieu. Peptide-MHC complexes (pMHCs) provide a census of the proteins being degraded by a cell at any given time. Cellular dysfunction, including viral infection or oncogenic transformation, alters the repertoire of presented peptide antigens as compared to those found during homeostasis. These peptides can then be recognized by T cells. In the case of class I MHCs, CD8+ cytotoxic T cells can recognize non-self pMHC antigens in a process that leads to killing of the targeted cell (Fig. 1A)^2^.

**Figure 1.**
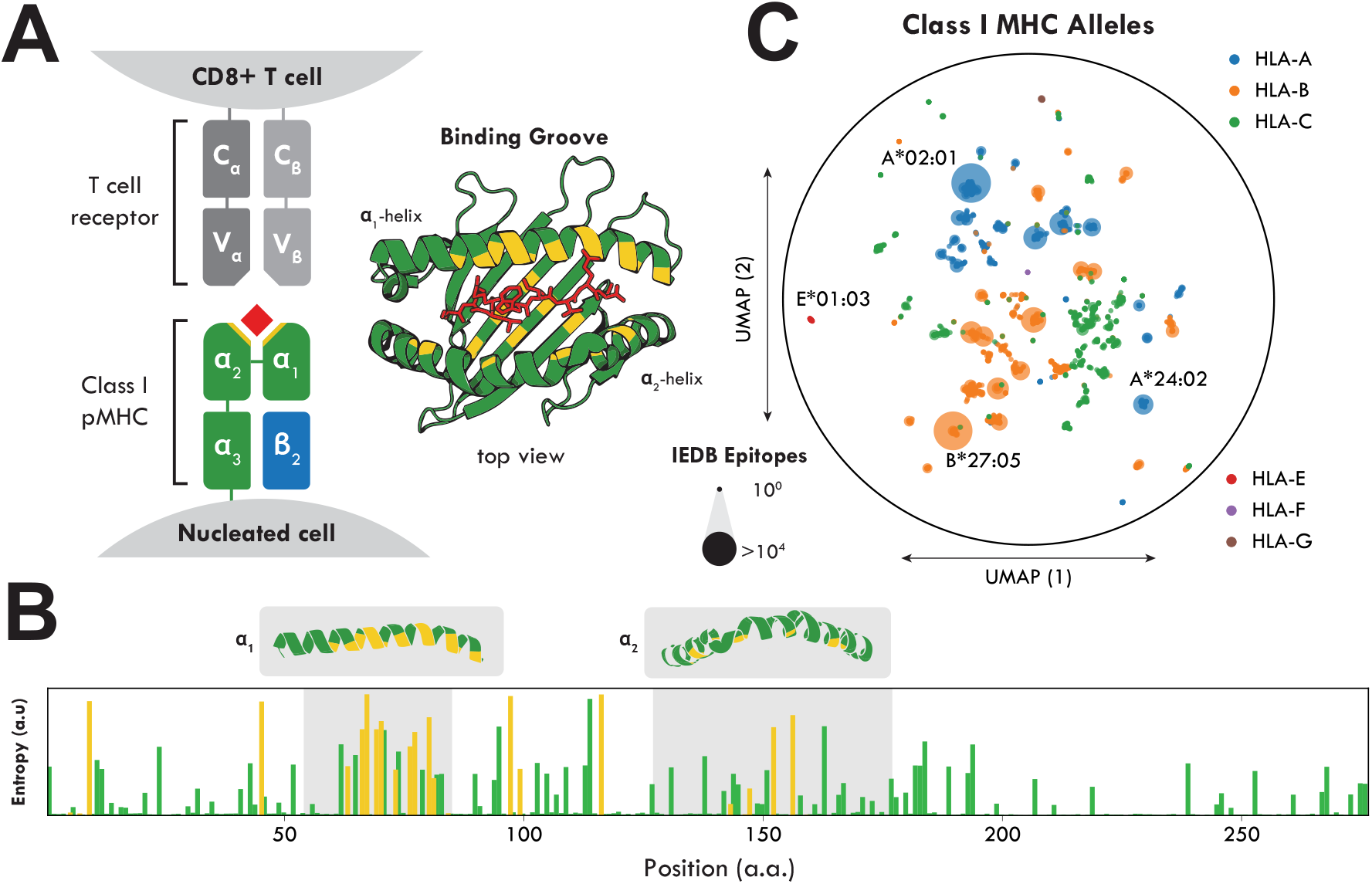
Class I MHC polymorphism. (A) MHCs contain a hydrophobic peptide binding groove that has numerous residues that directly bind the presented peptide. Here, we show a model peptide-MHC structure (NY-ESO-1157–165 in HLA-A2, PDB: 2BNR) where peptide contacts are highlighted in yellow. (B) MHCs carry significant polymorphism across all alleles. This polymorphism (measured as entropy in a global alignment of all class I MHC alleles) is specifically concentrated in peptide contacts, shown again in yellow. (C) For these thousands of distinct alleles, only a small fraction of this space has been thoroughly characterized for peptide binders. Each allele was projected into a 2D landscape based on protein sequence similarity. The size of each point corresponds to the number of known peptide binders found in the immune epitope database (IEDB). While a tremendous number of peptide binders are known for model alleles such as HLA-A*02:01 and B*27:05, many regions of this MHC sequence space are understudied.

The recognition of antigens by T cells uses multiple sources of immense molecular diversity to ensure effective protection from disease. MHCs are the most polymorphic genes in the genome, with thousands of coding variants across three class I MHC loci existing in the global population^3^. These polymorphisms cluster in the peptide-binding groove (Fig. 1B), which leads to each MHC having distinct preferences for the length and amino composition of bound antigens, especially at the “MHC anchor” positions of the peptide (typically P2 and PΩ for class I MHCs)^4^. Other factors, including cleavage preferences of the immunoproteasome^5^ and sequence similarity to proteins encoded in the host genome^6,7^, further affect which peptide antigens are recognized during an immune response.

The study of MHC antigen presentation and recognition has been aided by the development of a broad array of experimental and computational approaches. These methods have included X-ray crystallography^8–10^ and electron microscopy (EM)^11^ to understand the structural basis of antigen presentation, biophysical measurements of pMHC stability including fluorimetry^12^, and measurements of functional activity such as ELISPOT^13^. A number of high-throughput methods have recently emerged that allow detection of naturally occurring pMHCs using mass spectrometry^14–16^. Additionally, mammalian screening techniques have recently emerged using transduction of peptide variants^17^. Since even the highest throughput methods cannot experimentally analyze all possible peptides for a given MHC allele, computational tools have been developed to extrapolate patterns for peptide presentation for given alleles to new peptides^18–22^. While these approaches vary in algorithmic complexity, their performance are typically comparable, especially for the most commonly studied MHC alleles. This is likely because these methods rely on limited and often overlapping data to train their respective models^19,23,24^. Therefore, performance for a given allele is often correlated to the volume of existing data (Fig. 1C). Furthermore, existing training data can bias towards peptides derived from naturally occurring proteins^25^, hydrophilic peptides^26^, and peptides without cysteines^25^. This is largely due to there being relatively little training data, especially beyond the most studied MHC alleles.

Here, we describe a cost-effective, scalable platform for screening class I pMHC binding. MHCs are expressed on the surface of yeast with a random peptide. We have previously observed increases for β2M staining correlate with peptide binding for the given allele^27^. Through fluorescence-activated cell sorting (FACS) of pMHC libraries for β2M expression, we can identify the subset of peptides stabilizing the MHC via next-generation sequencing. Each selection can scan millions of random peptides, or hundreds of thousands of defined peptides by leveraging pooled oligo synthesis technology. We find this platform can additionally capture other biological features, such as the basis of HLA-B*57:01 abacavir hypersensitivity^28^. Ultimately, we apply this pipeline across sixteen class I alleles and identify antigen peptides derived from various infectious diseases (Mtb, SARS-CoV-2, Dengue, Zika). This platform not only identifies allele-specific peptide binders at scale but also generates new datasets across previously underexplored MHC alleles. By expanding empirical measurements of peptide binding, these results can feed back into computational models to improve prediction accuracy and deepen our understanding of allele-specific diversity in antigen presentation.

## Results

We and others have established yeast surface display as a high-throughput method to identify pMHC reactivity for a given antigen receptor^29–31^. In these methods, the peptide is typically covalently linked to the MHC via a Gly-Ser linker (Fig. 2A). Recently, we demonstrated that we can additionally use pMHC yeast display to identify peptide binding to class II MHCs independently of antigen receptor binding^25,32,33^. However, the method we utilized, cleaving the linker between peptide and -MHC linker in the presence of a soluble competitor peptide to identify encoded peptides that did not dissociate, was only feasible due to the open binding groove inherent to class II MHCs. For class I MHC yeast display, the closed peptide groove must be modified to allow the linker between the peptide and the MHC construct to proceed unimpeded^31,34,35^, which would not allow for a similar cleavage-based mechanism to measure peptide binding to MHC. Therefore, our previously described selection approaches do not extend to class I pMHC.

**Figure 2.**
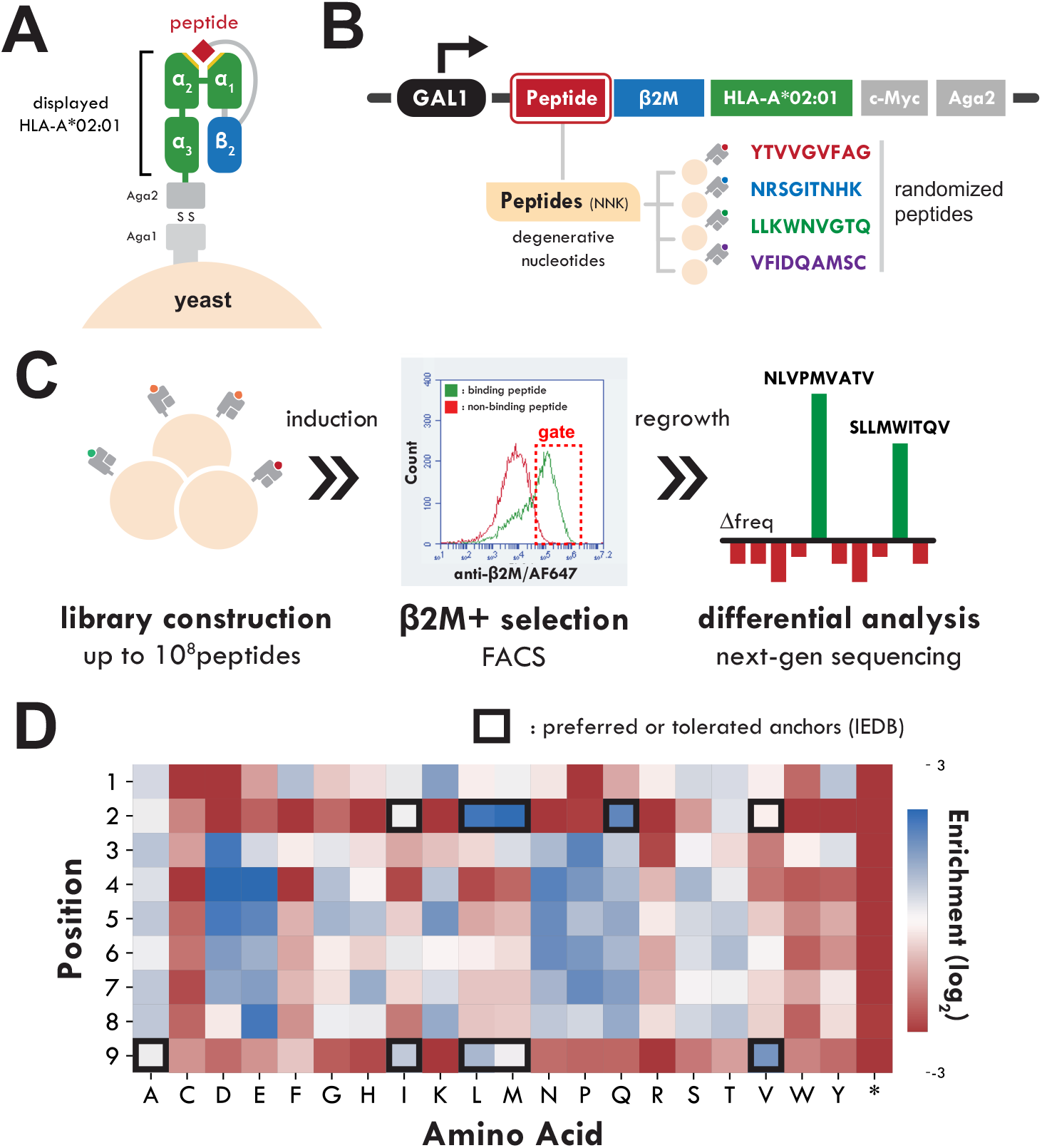
High-throughput class I pMHC screening pipeline. (A) In our screening method, a given class I MHC allele is expressed at the surface of yeast as a single chain trimer, linked to a randomized peptide at the N-termini. This construct is anchored to the surface via Aga2 to produce coat a single yeast cell. (B) Plasmids encoding this construct have their peptide randomized through degenerative nucleotides to give rise to recombinant libraries. (C) These mutagenized plasmids are electroporated in a pooled format, producing a library of yeast each transformed with a single peptide presented in a given MHC. Yeast with bound peptides help stabilize the surface complex and lead to higher pMHC expression. Using an anti-B2m antibody, this fraction of yeast can be isolated via fluorescence activated cell sorting (FACS). After 1-3 rounds of this enriching process, yeast populations can be harvested for plasmid and evaluated via next-generation sequencing for peptide context via differential analysis. (D) Pilot selection of peptides presented in HLA-A*02:01 produces an enrichment profile of amino acids in agreement with known allele-specific anchor residues (IEDB), demonstrating the feasibility of the selection pipeline to identify binding peptides in a randomize pool.

### A pMHC display platform for HLA-A*02:01 enables identification of binding peptides

To overcome this limitation, we extended our previous observations that class I pMHC expression on the surface of yeast correlates with peptide stability^34^ to hypothesize that high surface expression as measured by staining for β2M (an invariant subunit of class I MHCs) could be a means of selecting for class I MHC peptide binding motifs independently of binding to a cognate antigen receptor. To test this hypothesis, we generated random peptide libraries of 9 or 10 amino acids linked to HLA-A*02:01 displayed on yeast (Fig. 2B). These libraries were enriched via iterative rounds of selection for expression of β2M (Fig. 2C). Following two rounds of selection, each round of selection and starting libraries were sequenced to identify enriched peptide motifs. The fold change in frequency of each amino acid at position is represented in an enrichment map of the randomized peptide (Fig. 2D). Notably, the positions with the most distinct preferences are position 2 and position 9 which are known anchor residues of HLA-A*02:01. These results agree with the well-described amino acid preferences of positions 2 and 9^36^, suggesting our pipeline can successfully screen peptides capable of binding and stabilize class I pMHC.

### Deployment of display platform across twelve class I HLA alleles

To assess the generalizability of this approach, we expanded our screening platform to include 11 additional class I HLA alleles by modifying the plasmid backbone to contain the alternative MHCs (Fig. 3A). In addition to randomized peptides through degenerative nucleotides, we employed defined oligo pools to include sets of specific peptides with known affinity or binding for each allele, where available (Fig. 3B). These libraries were advanced in parallel through two rounds of our selection pipeline and evaluated for peptide enrichment via next-generation sequencing. In our randomized libraries, amino acid frequency was converted to an entropic representation of preferences by position (WebLogo^37^) (Fig. 3C). Across the twelve alleles, we find specific positions that carry high restriction for amino acids correlative of known anchor residues specific to alleles. Notably, many HLA-C alleles carry non-P2 anchor residues, which arise distinctly in each tested allele. As an additional validation, we utilized a database of known binding peptides from the Immune Epitope Database (IEDB)^38^ to validate the performance of our selections. Reported binding and non-binding peptides were evaluated for enrichment through selection (Fig. 3D). Each allele showed a distinct separation between these two groups, with known binding peptides presenting positive enrichment throughout selection. Finally, for a subset of class I HLA alleles we included peptides with known binding affinity (K_D_). Enrichment for these peptides is plotted against their reported affinity, where a strong correlation is seen across the four available alleles (Fig. 3E). Altogether, these results support the conclusion that our screening pipeline presents a robust solution to peptide-MHC binding.

**Figure 3.**
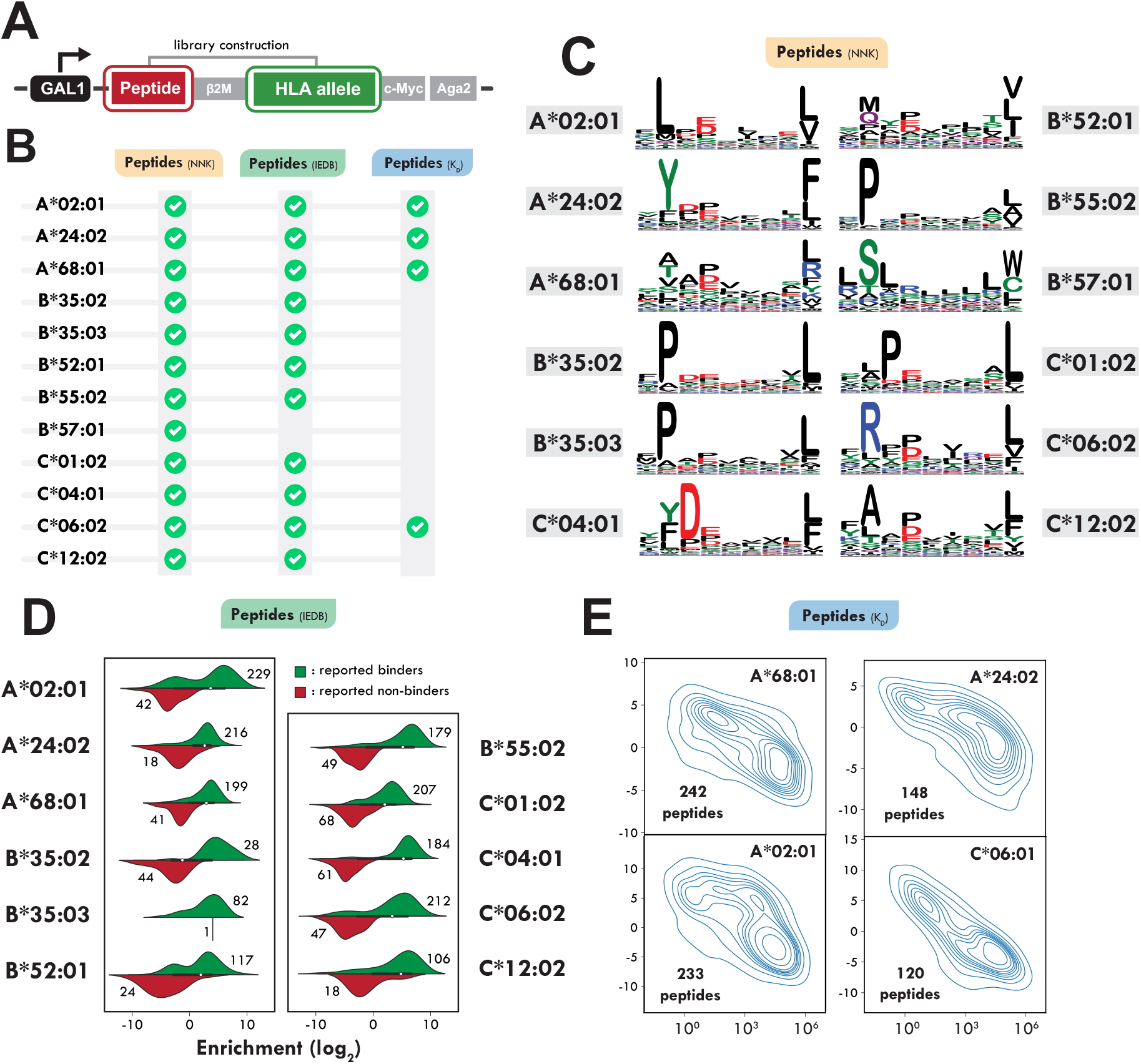
Validation across diverse range of MHC alleles. (A) The initial screening methodology was applied to a collection of class I MHC alleles to assess generalizability of the screening platform. In this iteration, the MHC backbone was replaced with new alleles, then adapted to the library generation step as previously described. (B) Each of these new HLA backbones were used to generate libraries with various sets of peptides, dependent on availability. These peptides could be fully randomized peptides (NNK), peptides previously reported as binding/non-binding (IEDB), or peptides with previously reported binding affinities (KD). (C) Randomized peptide libraries demonstrate a wide range of binding motifs distinct to each tested allele, as represented by sequence logos for enriched peptide fractions. (D) Oligo pool libraries for compatible alleles show distinct enrichment that is capable of differentiating binding and non-binding peptides at scale. (E) Enrichment score for peptides with known affinity correlate strongly with reported KD demonstrating an ability to resolve quantitative data.

### Recapitulation of MHC-mediated drug hypersensitivity

We sought to assess whether other elements of MHC biology could be recapitulated with our screening platform. A well-known drug hypersensitivity is the combination of patients carrying HLA-B*57:01 and treatment with abacavir, a small molecule commonly used to control HIV infection clinically. In previous research^19^, abacavir was shown to occupy the MHC binding grove, shifting the repertoire of peptides presented and inducing autoimmunity (Fig. 4A-B). To explore this, we cloned a randomized yeast library with HLA-B*57:01 and selected for peptide stability in induction media with abacavir included at concentrations ranging from 0 μg/mL to 500 μg/mL (Fig. 4C). After three rounds of FACS selection, a clear change in amino acid preferences is observed at P9, consistent with that of previous literature (Fig 4D). Additionally, individual peptide clones from the 0 μg/mL and 500 μg/mL selected libraries were induced with and without abacavir. We observed peptides selected without abacavir did not lose expression when induced with the small molecule, supporting the hypothesis the drug hypersensitivity expands the peptide repertoire rather than actively restricting existing peptides (Fig. 4E).

**Figure 4.**
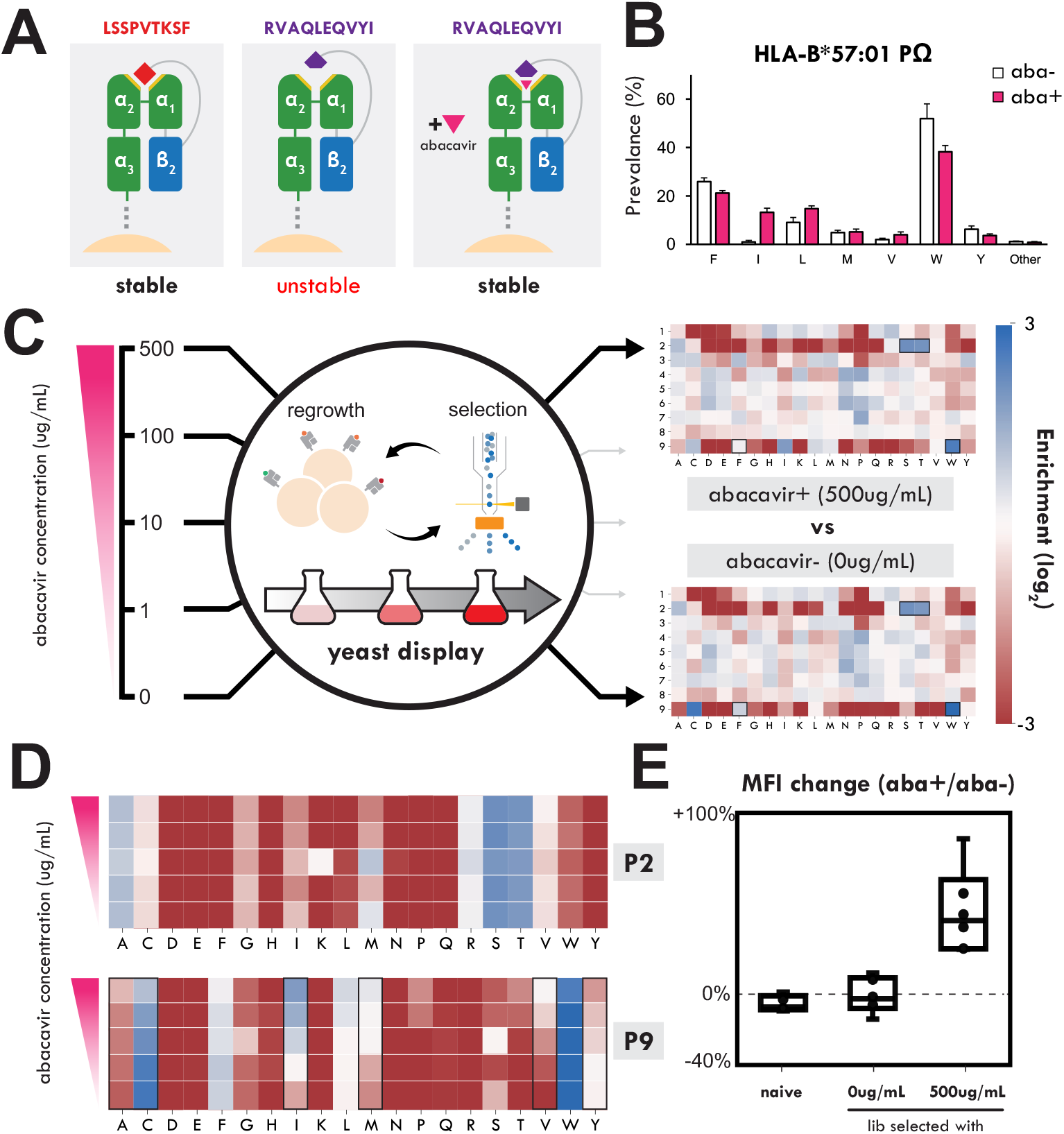
Resolving abacavir hypersensitivity in B*57:01. (A) HLA-B*57:01 carriers have been reported to have hypersensitivity to the small molecule abacavir. Previous studies have shown this relationship is achieved through abacavir stabilizing the binding groove of HLA-B*57:01 and expanding the repertoire of presentable peptides. (B) This extended repertoire primarily occurs on the final residue of the peptide, decreasing reliance on phenylalanine, F, and tryptophan, W, as seen in past mass spectrometry studies. (C) HLA-B*57:01 was implemented in our screening pipeline with randomized peptides induced in culture media spiked with varying concentrations of abacavir. The resulting selections produced enrichment maps of peptides specific to each tested abacavir concentration. (D) Position 2 in the enrichment map shows little variation in response to abacavir concentration, while position 9 shows expanded tolerance for non-canonical anchor residues such as isoleucine, methionine, and valine. (E) Libraries containing peptides selected in 500ug/mL of abacavir were uniquely able to express in the presence of the small molecule, in alignment with past observations.

### Rapid pathogen proteome screening

Finally, we aimed to use this platform to screen antigens from infectious diseases to both identify targets for immune responses in disease and potentially identify CD8+ T cell vaccine candidates. With this in mind, we constructed a defined peptide library that exhaustively screened pathogen-based peptides (Fig. 5A). First, reference genomes of major variants of SARS-CoV-2 at the time of the study, Type VII secretion products of tuberculosis (Mtb), all four serotypes of dengue, and the Zika virus proteome were gathered. Each protein/proteome was broken into 9mer and 10mer peptides via a sliding window and encoded in triplicate. The resulting oligo pool consisting of 594,351 sequences was then synthesized, amplified, and cloned into vectors for the twelve validated alleles plus an additional four alleles onboarded for this screen. These sixteen libraries were induced and sorted, which were separately grown and prepped for sequencing. Each of these libraries displayed enrichment motifs consistent with previous screens and known peptide motifs as well as passing an internal analytical step, suggesting successful enrichment of binding peptides.

**Figure 5.**
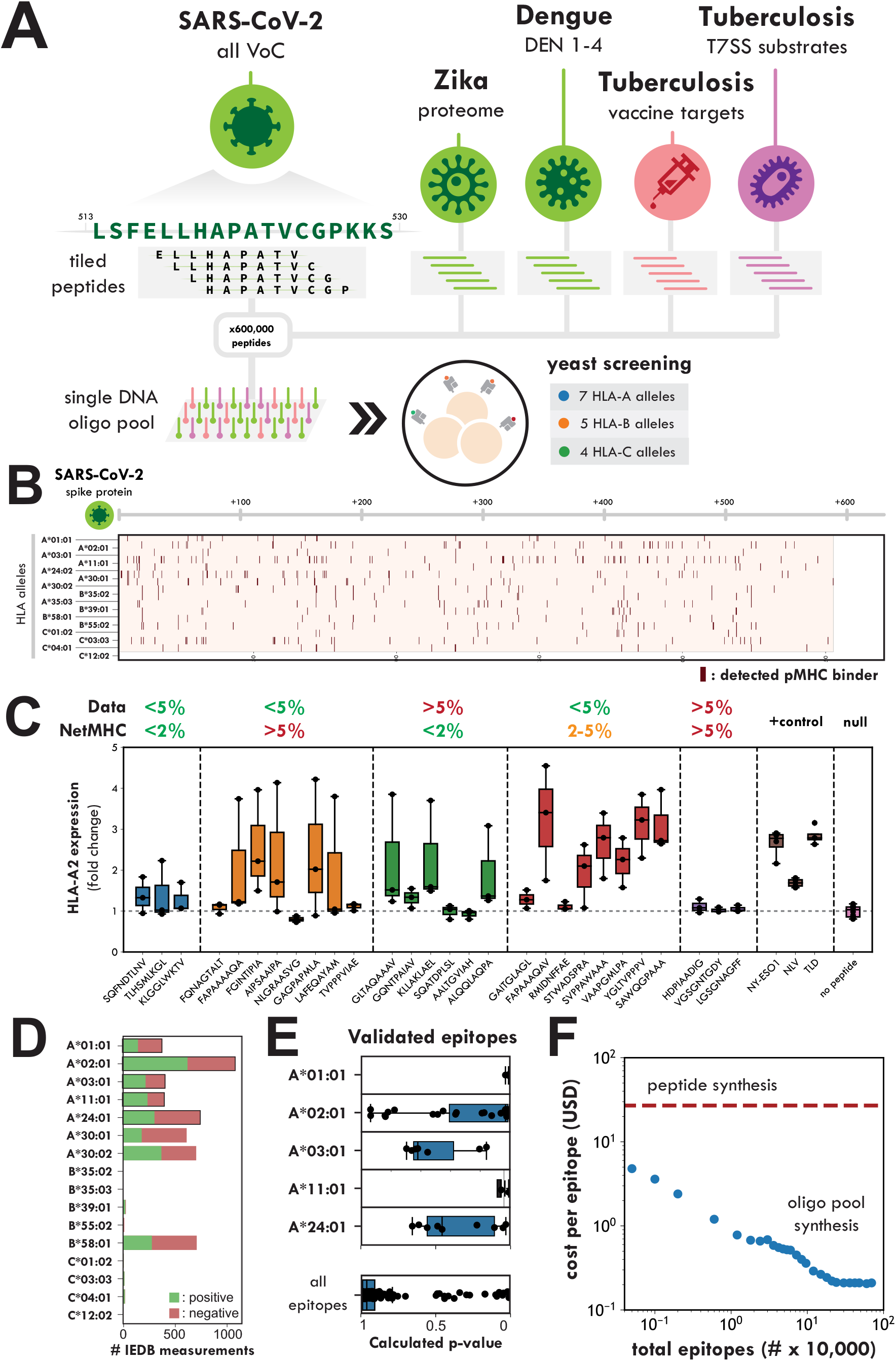
Rapid antigen screening using pMHC yeast display. Our platform enables the rapid identification of peptides capable of being presented as immune epitopes across a large number of class I alleles. (A) To this end, we selected four sets of proteins from various pathogens (Mtb T7SS substrates, Dengue, Zika, and SARS-CoV-2) to incorporate into a single defined oligo pool. These proteomes were fragmented into tiled peptides, and then added in triplicate using synonym codon encodings. Transformed libraries were run through two rounds of selection, then analyzed for peptide enrichment. (B) Using an analytic pipeline to bin peptides into binding & non-binding fractions, the binding peptides were mapped across all pathogen proteomes in unison. Shown is one resulting map for the SARS-CoV-2 spike protein across the sixteen successfully selected alleles. (C) Peptides from this screen were tested for binding in an HLA-A*02:01+ mammalian cell line in comparison to peptides predicted through NetMHC, a common computational peptide binding algorithm. (D) Among the sixteen tested alleles, the total number of reported peptides in the IEDB are shown highlighting the limited set of knowledge across most tested alleles. (E) Among these validated binding epitopes, p-values are shown for each peptide analyzed in the screening data showing some concordance. (F) Utilization of oligo pool synthesis demonstrates the tractability of screening large pools of peptides in parallel as compared to costs associated with individual peptide synthesis technologies.

We examined the location of binding peptides of various proteomes screened in our study across alleles. A representative map is shown in Fig. 5B, where we see the peptide binding map for SARS-CoV-2 spike protein across sixteen class I MHC alleles. Highlighted positions indicate the 9mer peptide at the given position was above the calculated cutoff for the given allele. To understand the relationship between these datasets and the computational prediction tools, NetMHC^37^, we randomly selected peptides with contrasting categorization as a peptide binder for A*02:01. This included peptides in agreement between NetMHC and internal datasets, peptides in disagreement between one of the two sources, and control peptides. Selected peptides were pulsed in A*02:01+ mammalian cell lines to resolve MHC stabilization via differential HLA expression (Fig. 5C). These results demonstrate the high potential for false negatives in NetMHC predictions (5/8 peptides <5% in yeast data ranking and >5% NetMHC ranking, 6/8 peptides <5% in yeast data ranking and 2-5% NetMHC ranking) as well as false negatives in the yeast data (4/6 peptides >5% in yeast data ranking and <2% NetMHC ranking).

Across the immune epitope database, a limited number MHC-ligand measurements have been deposited for peptides we have screened, with half of these alleles virtually unstudied (Fig. 5D). When considering specific epitopes with reported positive measurements in multiple studies, we found this screen to largely confirm stable peptide-MHC formation (Fig. 5E). Only a small fraction of peptides that our study reports, however, have been reported in these databases, pointing towards a lack of scalable tools capable of exploring pMHC binding. Existing tools rely upon arrayed peptide synthesis, which, for typical stability assays, costs $5-20 per peptide. In comparison, oligo pool synthesis enables the screening of peptides at much larger scales through synthesis costs that are mitigated by orders of magnitude (Fig. 5F).

## Discussion

We have applied pMHC yeast display to identify antigens presented by class I MHC alleles to improve understanding of allele-specific peptide presentation. Our data suggests that we can use peptide enrichment to recover peptide binding affinity (Fig 3C). In addition, even for alleles with large training datasets such as HLA-A*02:01, our comparisons with NetMHC predictions reveal mismatches between predicted non-binders and peptides consistently enriched in yeast display (Fig 5C). These findings support prior observations that NetMHC suffers from false negative predictions and highlight the utility of empirical peptide display platforms to uncover missed binders^39^. This approach to peptide binding is rapid and low-cost, providing an experimental basis to generate datasets that dramatically improve understanding of peptide presentation.

In this study, we establish antigenic maps of various pathogens (Mtb, SARS-CoV-2, Zika, Dengue) derived from a single oligo pool. The high throughput and relatively low cost of pooled oligonucleotide synthesis gives the ability to examine peptide-MHC interactions across many alleles at whole-proteome scale, providing a more complete view of possible presented antigens than previously possible. However, while this data provides a list of potential antigens, it is important to distinguish peptide immunogenicity from MHC binding. In disease, CD8+ T cell responses are restricted to the recognition of antigens presented by class I MHC alleles. Although peptide binding to class I MHCs is a prerequisite for a CD8+ T cell response, other factors also affect how likely it is that a peptide will be recognized by T cells. Antigenic protein expression level, stability, and proteolytic processing of the antigen in question affect whether it will be presented in living cells, and homology to self-proteins can affect the likelihood of a T cell response.

Despite the success of this platform, multiple considerations regarding the engineering of the pMHC molecules still exist. First, pMHC display uses native MHC sequences with a single engineered mutation, Y84A, which opens the F pocket of the MHC so peptides attached via a glycine-serine can be sterically accommodated^35^. While we observed PΩ residue preferences intact despite this change, it is possible biases are still imparted on our datasets. Additionally, the mechanism connecting β2M staining to stable peptide binding is unknown. We hypothesize these differences are driven by β2M accessibility as the class I MHC misfolds^40^. Interestingly, labeling conserved elements of class I MHCs via pan-allele antibodies was not an effective strategy for identifying MHC-binding peptides, potentially due to the numerous GS-linkers that interfere with epitope accessibility. Lastly, while the data presented here primarily focuses on 9mer selection data, class I MHCs can bind peptides with lengths ranging from 8-11 amino acids, with the exact distribution dependent upon the HLA allele. While many of our experiments included 8mers and 10mers, 9mer peptide binding motifs often conflated this data since any 10mer containing a valid 9mer register for the MHC at P1-P9 would still bind in the single chain trimer format. That is, if the PΩ residue in a 10mer does not sterically block binding, the peptide can shift into the 9mer register within the single chain trimer format. This data can be deconvoluted via computational measurements but provides additional analytical challenges.

Ultimately, this platform was applied successfully to sixteen distinct class I MHC alleles. There were nineteen other alleles that were screened that generated mild enrichments towards expected motifs, but with substantially reduced signal-to-noise. For these lower-performing alleles, additional FACS rounds or structural engineering strategies may help enhance peptide signal and expand the allele coverage of the platform. Interestingly, in our final proteomic screen, HLA-B alleles exhibited a 63% failure rate, markedly higher than HLA-A (13%) and HLA-C (17%). This may be driven by wild-type MHCs that fail to fold properly in yeast, or by peptide-free MHCs lacking sufficient destabilization to lower surface staining of β2M. Previous work with B*27:05 suggests the former, where stabilizing mutations are required before MHC display is able to enable TCR selection^41^. Despite these limitations, our platform offers a scalable, unbiased method to map peptide binding across diverse MHC alleles, laying groundwork for rapid antigen discovery and datasets to improve predictive modeling for class I MHC function via a platform that does not require the creation of MHC-specific reagents.

## Methods

### Construct design and allele selection

An initial A*02:01 containing plasmid was obtained from previous pMHC yeast display research^29,31^. Briefly, these plasmids encoded for the expression of an Aga2p fusion was placed under a GAL1 promoter. All vectors used the PLB1 signal peptide in place of the canonical AGA2 signal peptide to minimize peptide-specific expression bias as previously described^41,42^. Each element of these recombinant proteins (peptide, β2M, MHC, Myc epitope, Aga2p) were connected through GS-linkers. MHC protein sequences were acquired from IMGT and codon optimized for yeast using a custom script. Gene blocks were synthesized (Twist Biosciences), then replaced in the A*02:01 vector. Each vector was then confirmed via full plasmid sequencing (Plasmidsaurus).

Alleles were selected under multiple criteria. The first set of six alleles (A*02:01, A*24:02, A*68:01, B*35:02, B*35:03) were already cloned with an ongoing clinical collaboration. The next set of alleles (eleven new alleles, see supplemental materials) were selected due to known disease associations ^42,43^. In principle, further study of these alleles could contribute to why these alleles correlate with disease. The final set of eighteen alleles were selected under multiple conditions. First, one set was selected for their prevalence in Mtb-high geographic regions. Another set of alleles was selected as the MHCs present in a cell line often used for Mtb co-culture experiments (THP-1). Lastly, a set of alleles were selected as a set of globally abundant alleles.

### Clonal yeast selections

To explore differential staining for each allele, multiple clonal yeast populations were created that contained a single peptide linked to a given MHC. For each allele, this included positive, known binders, as well as an irrelevant peptide (non-binder). Binding peptides were selected from the IEDB as the top three most cited peptides for each allele^43^. Non-binding peptides were selected randomly from binding peptides for other alleles (list of peptides are recorded in the supplemental materials). In the pathogen library screens, the non-binding peptide was uniformly chosen as a 9mer composed of only alanine. For each of these peptides, oligos were ordered (IDT) and amplified using primers containing homology to each vector. BsaI digested vector was mixed with these products and assembled using Gibson cloning reaction mix (Genscript). Plasmids were confirmed through Sangar sequence (Azenta Biosciences), then transformed and plated using competent yeast (Frozen-EZ yeast transformation kit, Zymo Research). A single colony was picked and cultured in SD-CAA, then induced in SG-CAA for 48 hours before characterization via flow cytometry (Accuri, BD).

### Library construction and electroporation

Each vector containing flanking BsaI restriction enzyme cut sites across the peptide-containing region of the plasmid. Purified plasmid was incubated with BsaI-HFv2 (NEB) overnight and purified via gel extraction. Separately, either degenerative oligos (IDT) or oligo pools (Twist) were amplified with primers that contained homology to the backbone for all vectors. PCR products were purified using SPRIselect beads (x1.5 ratio, Beckman Coulter). Between 2-5ug of digested vector was combined with a 20x excess of purified insert, and was electroporated using a protocol analogous to Deventer et al^44^. All libraries produced in this study had diversity at least 20x over the oligo pool diversity, maximizing the likelihood all designed constructs would be present in the library.

### FACS sorting

Each library was stored at 4°C until usage, where they were grown in SD-CAA media overnight. Yeast were then resuspended in SG-CAA media (OD: 1.0) to be induced at 20°C for 48 hours. After this, yeast were spun down, washed with FACS buffer, and stained with anti-β2M Alexa 647 antibody (AbCam) at 200:1 dilution for 20 minutes. Cells were again washed, then resuspended at a density of approximately 50 million/mL and run on FACS (FACS Aria II, BD). Gates were drawn at a slant (compensating with side scatter) in which the highest MFI fractions would be selectively sorted (top 1%/5% in NNK libraries, top 1%/5% in IEDB libraries, top 1%/1% in pathogen libraries). Each library was sorted at approximately 20x the diversity of each oligo pool over 20-40 minutes per library. Cells were recovered in SD-CAA and grown until confluent. For selections involving abacavir, abacavir was spiked into the induction medium (SG-CAA), ranging from 0ug/mL to 500ug/mL.

### Library preparation and sequencing

Recovered yeast populations (post-selection) and induced populations (pre-selection) were harvested for plasmid recovery and sequencing. To do this, 20-50 million yeast were pelleted for plasmid recovery using a zymoprep yeast plasmid miniprep (Zymo Research). Half of the plasmid elution was mixed with a PCR mix (KAPA HiFi HotStart ReadyMix, Roche) along with primers containing Illumina Truseq primer handles. The PCR products were purified using SPRIselect beads (x1.2 ratio, Beckman Coulter) and analytically evaluated via DNA gel. Afterward, beads were amplified using a custom primer set that contained unique dual indices and the p5/p7 Illumina handles. Samples were then run on an Illumina Miseq (NNK libraries) or an Illumina NextSeq (IEDB and pathogen libraries). Up to 96 samples were multiplexed in a single sequencing run.

### Data analysis

Data analysis was performed using custom scripts using methods common to past yeast sequencing pipelines^45^. Briefly, reads were demultiplexed by sample index and filtered for 5’ and 3’ homology. Each read was mapped to the nearest matching oligo (hamming distance of 1) or simply stored (in the case of the NNK libraries). Read counts for each peptide sequence for each screening campaign are included in the supplementary materials. Multiple encodings of a single peptide were then summed prior to calculating the enrichments for each unique peptide between selected and naïve populations using:

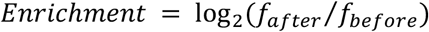

For heatmap generation, each oligo was translated into protein space, and split by peptide size. Subsequently, count matrices were generated for each sample and subpopulation that recorded the number of occurrences of a given residue at a given position in the peptide. Analyses were all performed in Python 3.7, and figures were generated using matplotlib^46^ and seaborn^47^ packages, with individual scripts available upon request.

### Binning algorithm for pathogen peptides selections

To call peptide binders in the pathogen screen for each allele, peptides were first ordered by read frequency in the selected population. The binding motif, with relative enrichments for each amino acid at each position, was used to score each of these peptides for how likely this peptide is a part of the signal. This was calculated using:

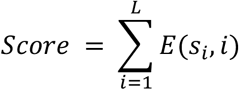

where *E*(*s, i*) is the log_2_ enrichment of an amino acid *s* at position *i, s*_*i*_ is the residue of the given peptide at position *i*, and *L* is the length of the peptide. The top ranked peptides for selection score highly in this metric, while low ranked peptides score poorly (indicating noise). We calculated a rolling average of this metric (*n* = 1000) and considered the transition from signal to noise to occur at the 25^th^ percentile. Plots for each allele are shown in the supplemental materials. For peptides lying above this threshold, we called them a binding peptide for that given allele. With this classifier, we generated protein maps for all pathogen proteins included in the study using custom scripts. To compare the performance of our screen against existing prediction tools, we ran NetMHCpan4.1b^17^ on all pathogen peptides and reported the eluted ligand rank (included in supplemental materials).

### Peptide pulse in A*02:01+ mammalian cell lines

To validate peptide binding in a mammalian context, T2 cells (A*02:01+) were pulsed with peptides and co-cultured with TCR-expressing T cells. Briefly, T2 were plated in a U-bottom 96-well plate in 50uL RPMI 1640 supplemented with 10% FBS and incubated at 37°C. Crude peptides were synthesized (Genscript) and diluted to a 200 μM working solution in RPMI. After peptide dilution, 50 μL of each peptide solution was added to the corresponding wells containing T2 cells, mixed gently, and incubated for 24 hours at 37°C. Cells were then stained with fluorophore-conjugated antibodies against HLA-A2 and analyzed on flow cytometry to measure increased expression of HLA-A2 via peptide-mediated stabilization of the binding groove.

## Data Availability

All raw and processed data supporting the findings of this study, including read counts, enrichment values, peptide classifications, NetMHC predictions, and reference sequences, are provided in the Supplementary Data Directory with detailed file descriptions. Raw sequencing files have been deposited in the NIH Sequence Read Archive under accession number SUB15276783. All additional data and analysis scripts are available from the corresponding author upon request.

## Acknowledgements

We thank the Koch Institute’s Robert A. Swanson (1969) Biotechnology Center for their technical support, especially the Flow Cytometry Facility and MIT BioMicro Center. This work was supported by the National Institutes of Health (DP2-AI158126 and R21-AI156664 to M.E.B., and R01-AI022553 and R35-GM142900 to B.D.B.), the Packard Foundation, and Schmidt Futures for M.E.B., and a National Science Foundation Graduate Fellowship to P.V.H. This work was additionally supported in part by the Koch Institute Support (core) Grant P30-CA14051 from the National Cancer Institute.

## Supplemental Information

Supplemental information for this manuscript can currently be found at: http://bit.ly/3H8C1ge

